## Supplemental for "Syngeneic natural killer cell therapy activates dendritic and T cells in metastatic lungs and effectively treats low-burden metastases"

#### Supplemental Figures

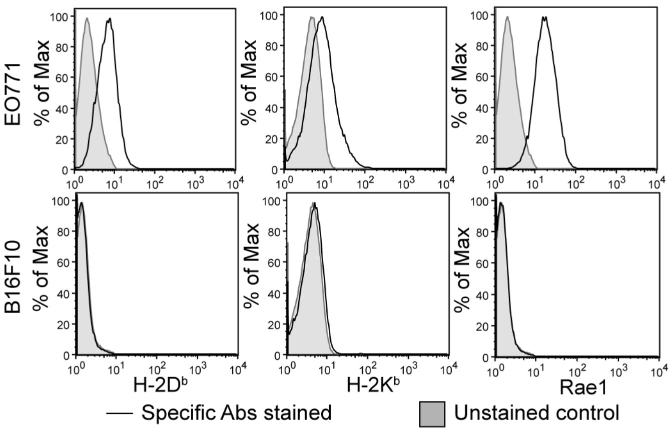


**Figure 1-figure supplement 1.** Expression of MHC-I molecules and Rae-1 by EO771 and B16F10 cells *in vitro* **(**related to Figure 1E)**.**

Representative flow plots of MHC-I molecules and Rae-1 expression by EO771 and B16F10 cells from three independent experiments are shown.

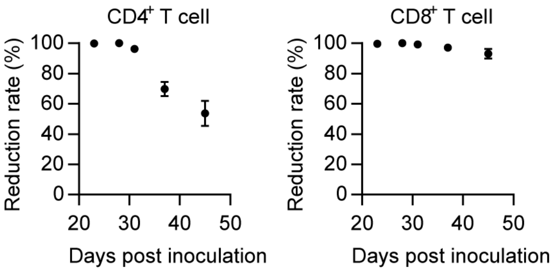


**Figure 3-figure supplement 1.** Effectiveness of *in vivo* depletion of $CD4^+$ and $CD8^+$ T cells by antibody (related to Figure 3D).

Mice were treated with CD4- or/and CD8α-specific antibodies at day 19 post tumor inoculation, and then underwent tumor resection at day 21, followed by NK cell therapy during days 24-31. Levels of circulating $CD4^+TCR\beta^+$ cells in the blood were reduced by 96-100% during the period of NK cell transfer, and by 70% and 54%, respectively, at 1- and 2-weeks post NK cell transfer. The level of blood-circulating $CD8^+TCR\beta^+$ cells was reduced by 99-100% during the period of NK cell transfer, and by 97% and 93%, respectively, at 1- and 2-weeks post NK cell transfer. Each time point compiles data from 3-10 mice from three independent experiments and is shown as mean ± SEM.

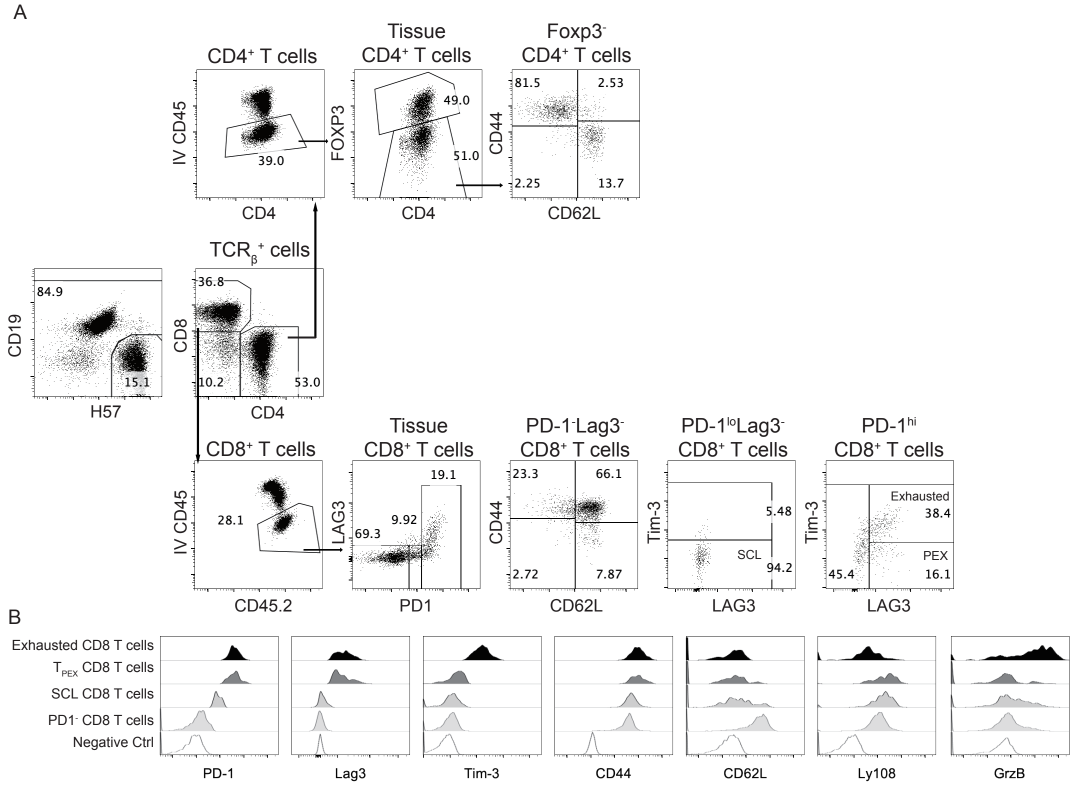


**Figure 5-figure supplement 1.** Analysis of T cells in lung tissue.

(A) Representative flow plots show the gating of CD4$^+$ and CD8$^+$ T cell subsets. (B) Histograms show the expression of indicated molecules that mark the four CD8$^+$ T cell subsets from a representative NK-cell-treated mouse. The negative controls are either FMO or single stain of a different molecule.

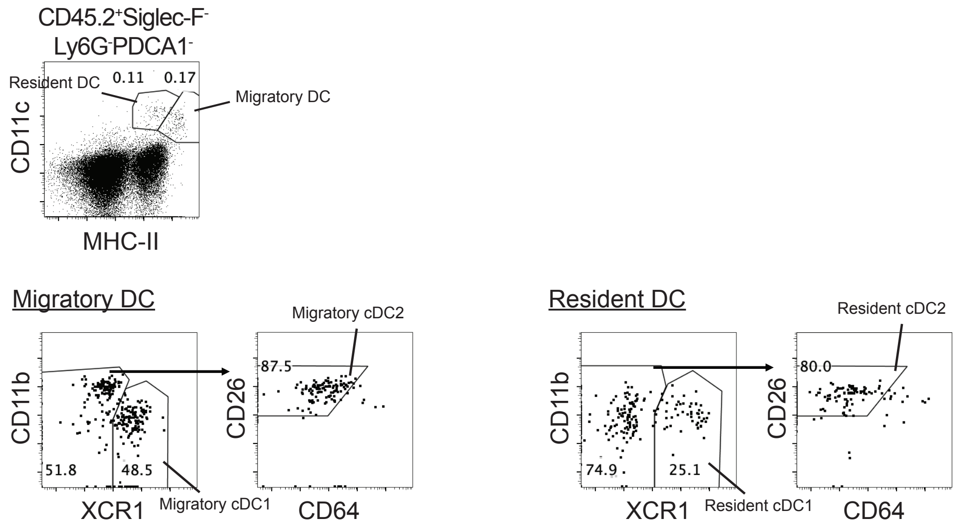


**Figure 6-figure supplement 1.** Analysis of cDCs in mLN.

Representative flow plots show the gating of migratory and resident cDCs in mLN.

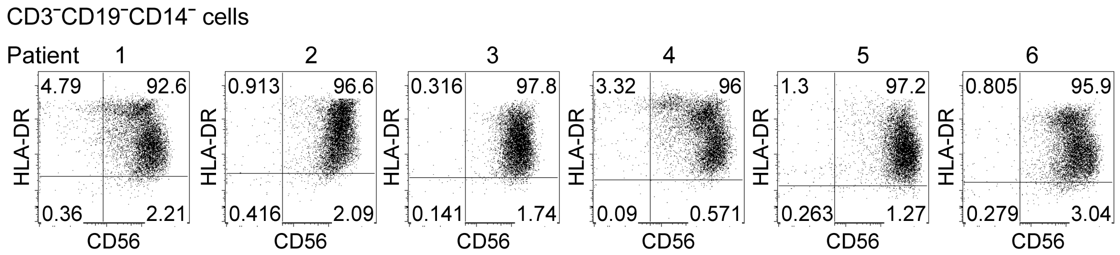


**Figure 8-figure supplement 1.** Expression of CD56 and HLA-DR by CD3$^-$CD19$^-$CD14$^-$ cells after *ex vivo* expansion (related to Figure 8A)**.**

Flow plots representative of the six batches of cell preparation for each patient are shown.

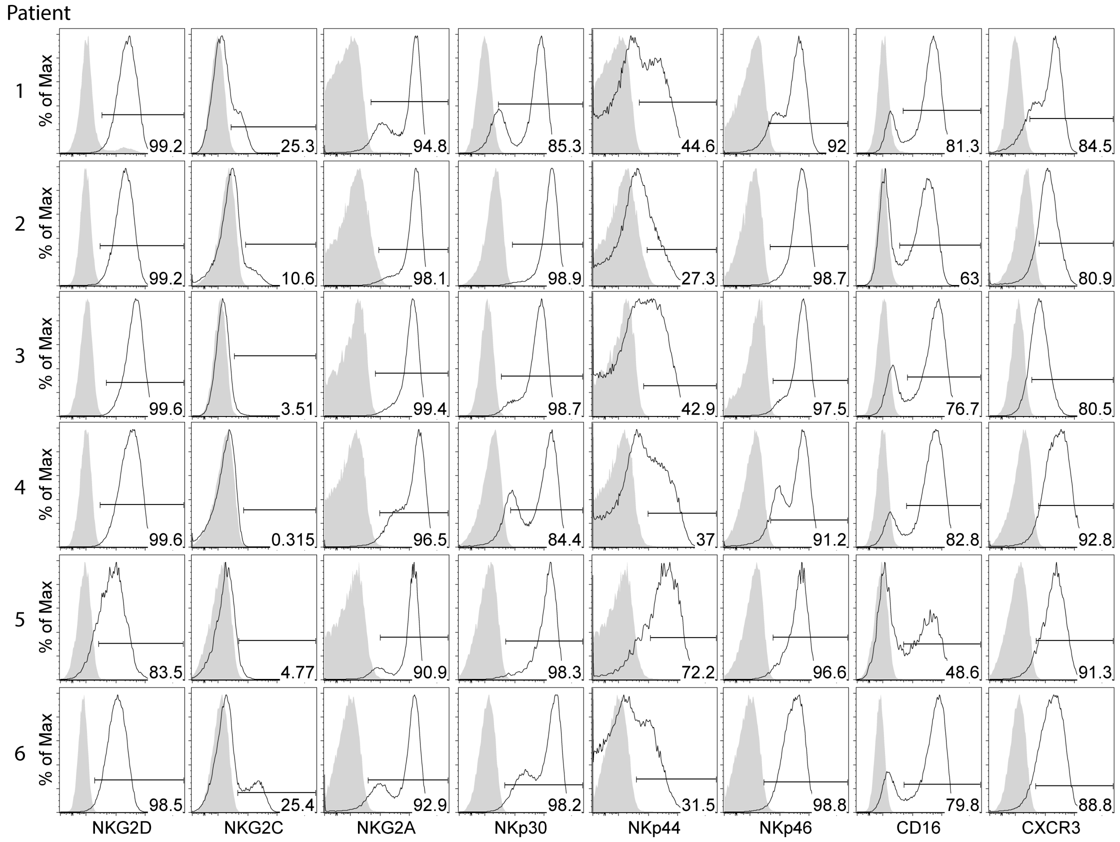


**Figure 8-figure supplement 2.** Expression of NKRs, CD16 and CXCR3 by the expanded HLA-DR+ NK cells (related to Figure 8B).

Flow histograms representative of the six batches of cell preparation for each patient are shown. The solid line and the filled gray peak represent gated HLA-DR+ NK cells stained with and without specific antibody, respectively.

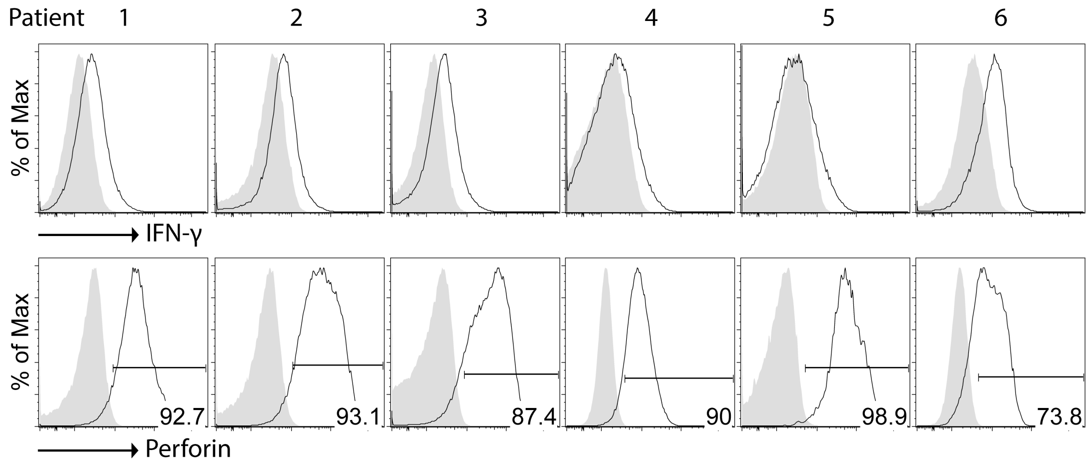


**Figure 8-figure supplement 3.** Expression of IFN-γ and perforin in the expanded HLA-DR$^+$ NK cells (related to Figure 8C).

Flow histograms representative of the six batches of cell preparation for each patient are shown. The solid line and the filled gray peak represent gated HLA-DR$^+$ NK cells stained intracellularly with specific antibody and isotype control antibody, respectively.

### Supplemental Materials

**Antibodies used for FACS analysis of murine cells**

| Species | Protein | Vendor | Catalog Number | Clone |
|---|---|---|---|---|
| mouse | B220 | BioLegend | 103224 | RA3-6B2 |
| | CCR5 | eBioscience | 12-1951-82 | HM-CCR5(7A4) |
| | CCR7 | BioLegend | 120106 | 4B12 |
| | CCR7 | BioLegend | 120109 | 4B12 |
| | CD11b | BD | 741934 | M1/70 |
| | CD11b | BioLegend | 101228 | M1/70 |
| | CD11c | BioLegend | 117316 | N418 |
| | CD11c | BioLegend | 117308 | N418 |
| | CD11c | eBioscience | 63-0114-82 | N418 |
| | CD11c | eBioscience | 64-0114-82 | N418 |
| | CD19 | BioLegend | 115578 | 6D5 |
| | CD19 | BioLegend | 115506 | 6D5 |
| | CD19 | BioLegend | 115528 | 6D5 |
| | CD19 | eBioscience | 366-0193-82 | 1D3 |
| | CD26 | BD | 741492 | H194-112 |
| | CD27 | BioLegend | 124216 | LG.3A10 |
| | CD4 | BioLegend | 100480 | GK1.5 |
| | CD40 | BioLegend | 124622 | Mar-23 |
| | CD44 | BD | 741227 | IM7 |
| | CD44 | BioLegend | 103044 | IM7 |
| | CD45 | BioLegend | 103108 | 30-F11 |
| | CD45.2 | BD | 564616 | 104 |
| | CD62L | BD | 740660 | MEL-14 |
| | CD63 | BioLegend | 143922 | NVG-2 |
| | CD64 | BioLegend | 139320 | X54-5/7.1 |
| | CD64 | BioLegend | 139304 | X54-5/7.1 |
| | CD80 | eBioscience | 15-0801-82 | 16-10A1 |
| | CD86 | BD | 741737 | GL1 |
| | CD86 | BioLegend | 105042 | GL-1 |
| | CD8a | BioLegend | 100740 | 53-6.7 |
| | CD8a | eBioscience | 365-0081-82 | 53-6.7 |
| | CD8a | eBioscience | 64-0081-82 | 53-6.7 |
| | CXCR3 | eBioscience | 12-1831-82 | CXCR3-173 |

| | | | | |
|---|---|---|---|---|
| | CXCR6 | BioLegend | 151103 | SA051D1 |
| | DNAM-1 | BioLegend | 128806 | 1.00E+06 |
| | EOMES | eBioscience | 46-4877-42 | Dan11mag |
| | Foxp3 | eBioscience | 15-5773-82 | FJK-16s |
| | Granzyme B | BioLegend | 515408 | GB11 |
| | Granzyme B | BioLegend | 372208 | QA16A02 |
| | H-2 Class I | BD | 749712 | M1/42 |
| | H-2Db | BioLegend | 111508 | KH95 |
| | H-2Kb | eBioscience | 17-5958-82 | AF6-88.5.5.3 |
| | H-2Kb/H2Db | BioLegend | 114612 | 2006/8/28 |
| | I-A/I-E | BD | 750281 | M5/114.15.2 |
| | I-A/I-E | BD | 751570 | M5/114.15.2 |
| | I-A/I-E | BioLegend | 107626 | M5/114.15.2 |
| | IFN-g | eBioscience | 367-7311-82 | XMG1.2 |
| | IFN-g | BioLegend | 505808 | XMG1.2 |
| | IL-12/23p40 | BioLegend | 505206 | C15.6 |
| | Ki67 | BioLegend | 652469 | 16A8 |
| | Lag-3 | BioLegend | 125219 | C9B7W |
| | Ly108 | eBioscience | 741679 | 13G3 |
| | Ly49A | BioLegend | 116805 | YEL/48.10.6 |
| | Ly49D | BioLegend | 138303 | 4.00E+05 |
| | Ly49G2 | eBioscience | 11-5781-82 | eBio4D11 |
| | Ly49H | BioLegend | 144712 | 3D10 |
| | Ly49I | eBioscience | 11-5895-82 | YLI-90 |
| | Ly6C | BioLegend | 128028 | HK1.4 |
| | Ly6G | BioLegend | 127645 | 1A8 |
| | Mouse IgG1, κ | BioLegend | 400151 | MOPC-21 |
| | Mouse IgG1, κ | eBioscience | 46-4714-82 | P3.6.2.8.1 |
| | Mouse IgG1, κ | eBioscience | 50-4714-82 | P3.6.2.8.1 |
| | NK1.1 | BioLegend | 108714 | PK136 |
| | NK1.1 | BioLegend | 108728 | PK136 |
| | NK1.1 | eBioscience | 364-5941-82 | PK136 |
| | NKG2A | eBioscience | 46-5897-82 | 16A11 |
| | NKG2D | eBioscience | 12-5882-82 | CX5 |
| | PD-1 | eBioscience | 48-9981-82 | RMP1-30 |
| | PDCA1 | BioLegend | 127038 | 927 |

| | | | | |
|---|---|---|---|---|
| | PDCA1 | BioLegend | 127019 | 927 |
| | PD-L1 | eBioscience | 63-5982-82 | MIH5 |
| | Rae-1 | R&D | FAB17582P | 186107 |
| | Rat IgG 2a, κ | eBioscience | 15-4321-82 | eBR2a |
| | Rat IgG1, κ | BioLegend | 400412 | RTK2071 |
| | Rat IgG1, κ | BioLegend | 400408 | RTK2071 |
| | Rat IgG1, κ | eBioscience | 367-4301-81 | eBRG1 |
| | siglecF | BD | 747316 | E50-2440 |
| | siglecF | BD | 740158 | E50-2440 |
| | siglecF | eBioscience | 78-1702-82 | 1RNM44N |
| | T-bet | eBioscience | 50-5825-82 | eBio4B10 |
| | TCRb | BD | 749915 | H57-597 |
| | TCRb | BioLegend | 109240 | H57-597 |
| | TCRb | homemade | | H57-597 |
| | TCRd | homemade | | GL3 |
| | Tim-3 | BioLegend | 119716 | RMT3-23 |
| | TRAIL | BioLegend | 109306 | N2B2 |
| | XCR1 | BioLegend | 148216 | ZET |
| | XCR1 | BioLegend | 148225 | ZET |

**Antibodies used for FACS analysis of human cells**

| Species | Protein | Vendor | Catalog Number | Clone |
|---|---|---|---|---|
| Human | CD14 | BioLegend | 301820 | M5E2 |
| | CD159 | Beckman Coulter | IM3291U | Z199 |
| | CD16 | eBioscience | 48-0168-42 | eBioCB16 |
| | CD183 | BioLegend | 353712 | G025H7 |
| | CD19 | eBioscience | 47-0199-42 | HIB19 |
| | CD3 | BioLegend | 300426 | UCHT1 |
| | CD3 | eBioscience | 48-0038-42 | UCHT1 |
| | CD314 | BioLegend | 320822 | 1D11 |
| | CD335 | BioLegend | 331908 | 9.00E+02 |
| | CD336 | BioLegend | 325108 | P44-8 |
| | CD337 | BioLegend | 325212 | P30-15 |
| | CD56 | Invitrogen | 25-0567-42 | CMSSB |
| | HLA-DR | BioLegend | 307632 | L243 |
| | HLA-DR | eBioscience | 48-9952-42 | L243 |

| | | | | |
|---|---|---|---|---|
| | IFN-γ | BioLegend | 502516 | 4S.B3 |
| | NKG2C | R&D | FAB138A | 134591 |
| | Perforin | eBioscience | 12-9994-42 | dG9 (delta G9) |

**Sequences of primers used for qRT-PCR**

| Gene name | Forward primer | Reverse primer |
|---|---|---|
| Cxcl9 | GGAGTTCGAGGAACCCTAGTG | GGGATTTGTAGTGGATCGTGC |
| Cxcl10 | CCAAGTGCTGCCGTCATTTTC | GGCTCGCAGGGATGATTTCAA |
| Cxcl11 | GGCTTCCTTATGTTCAAACAGGG | GCCGTTACTCGGGTAAATTACA |
| Ccl3 | TTCTCTGTACCATGACACTCTGC | CGTGGAATCTTCCGGCTGTAG |
| Ccl4 | TTCCTGCTGTTTCTCTTACACCT | CTGTCTGCCTCTTTTGGTCAG |
| Ccl5 | GCTGCTTTGCCTACCTCTCC | TCGAGTGACAAACACGACTGC |
| Cxcl16 | CCTTGTCTCTTGCGTTCTTCC | TCCAAAGTACCCTGCGGTATC |
| Cypa | GAGCTGTTTGCAGACAAAGTTC | CCCTGGCACATGAATCCTGG |
